# DdmDE eliminates plasmid invasion by DNA-guided DNA targeting

**DOI:** 10.1101/2024.07.20.604412

**Authors:** Xiao-Yuan Yang, Zhangfei Shen, Chen Wang, Kotaro Nakanishi, Tian-Min Fu

## Abstract

Horizontal gene transfer is a key driver of bacterial evolution, but it also presents severe risks to bacteria by introducing invasive mobile genetic elements. To counter these threats, bacteria have developed various defense systems, including prokaryotic Argonautes (pAgo) and the <u>D</u>NA <u>D</u>efense <u>M</u>odule DdmDE system. Through biochemical analysis, structural determination, and *in vivo* plasmid clearance assays, we elucidate the assembly and activation mechanisms of DdmDE, which eliminates small, multicopy plasmids. We demonstrate that DdmE, a pAgo-like protein, acts as a catalytically inactive, DNA-guided, DNA-targeting defense module. In the presence of guide DNA, DdmE targets plasmids and recruits a dimeric DdmD, which contains nuclease and helicase domains. Upon binding to DNA substrates, DdmD transitions from an autoinhibited dimer to an active monomer, which then translocates along and cleaves the plasmids. Together, our findings reveal the intricate mechanisms underlying DdmDE-mediated plasmid clearance, offering fundamental insights into bacterial defense systems against plasmid invasions.

## INTRODUCTION

Horizontal gene transfer (HGT), also known as lateral gene transfer, involves the exchange of genetic material among distantly related organisms, shaping evolution across diverse life forms^1,2^. In bacteria, HGT is a primary driver of antibiotic resistance but also poses significant risks, as mobile genetic elements like integrative plasmids can disrupt genomic integrity ^3^. In response, bacteria have evolved diverse defense systems^4–6^. Short Argonaute (Ago) systems, such as SPARTA, offer effective immunity against plasmid invasion by depleting NAD^+^, inducing bacterial suicide for community protection^7,8^. Additionally, *Clostridium butyricum* Ago (*Cb*Ago) was shown to be a bacterial immune defense module that can suppress plasmid propagation and phage infection ^9^. The <u>D</u>NA <u>D</u>efense <u>M</u>odule DdmDE, discovered in *Vibrio cholerae* strain 7PET, represents another anti-plasmid defense system with an unclear mechanism^10^.

The DdmDE anti-plasmid system was discovered in a *Vibrio cholerae* strain of the O1 EI Tor biotype (7PET), which is the primary cause of the seventh pandemic. From a representative 7PET strain A1552, two plasmid defense systems were identified and were named as DdmABC and DdmDE, respectively ^10^. DdmABC can defend against both plasmid invasion and bacteriophage infection^10^. In contrast, DdmDE is mainly responsible for eliminating plasmids, particularly small, multi-copy plasmids, by degradation^10^. Sequence analysis revealed that DdmD is composed of a superfamily II (SF2) helicase domain and a PD-(D/E)XK nuclease domain, while DdmE resembles an Argonaute protein with a MID domain and a PIWI-like domain^11–15^. Both DdmD and DdmE are required for bacterial anti-plasmid defense, yet their cooperative degradation mechanism remains elusive^10^.

Here, we integrate biochemical analysis, structural determination, and *in vivo* assays to unveil a captivating mechanism that governs the assembly and activation of the DdmDE anti-plasmid defense system. We reveal that DdmE encompasses all the characteristic domains found in Argonautes, including N, L1, PAZ, L2, MID, and PIWI, along with a domain inserted within the PIWI domain to facilitate interaction with DdmD^16^. The cryo-EM structure of the DdmE-DNA duplex complex, combined with biochemical assays, demonstrates that DdmE functions as a catalytically inactive, DNA-guided, DNA-targeting defense module. Notably, unlike typical monomeric SF2 helicases^17–21^, DdmD exists as a dimer in its apo state. Upon binding to DNA substrates, the positively charged channel of DdmD accommodates DNA, triggering disassembly of the dimeric form. Subsequently, the monomeric DdmD, propelled by its helicase domain, translocates along the plasmid DNA and efficiently eradicates plasmid DNA via its nuclease domain.

Our study provides a comprehensive mechanistic understanding of DdmDE as a DNA-guided, DNA-targeting anti-plasmid defense system, laying the foundation for harnessing the system as a molecular tool in biotechnology.

## RESULTS

### Apo DdmD is an autoinhibited dimer

To elucidate the assembly of DdmD, we determined its cryo-EM structure to a resolution of 3.1 Å using single-particle analysis (Figures S1A-S1C, and Table S1). Unlike the typical monomeric assembly of other known SF2 helicases^22^, apo DdmD forms a C2 symmetric homodimer with dimensions of 115 Å ξ110 Å ξ 65 Å (Figures 1A and 1B). Each protomer consists of five domains: helicase domain 1 (HD1), the inserted domain for dimerization (IDD), the Arch domain, helicase domain 2 (HD2), and the nuclease domain (Figures 1C and 1D).

**Figure 1.**
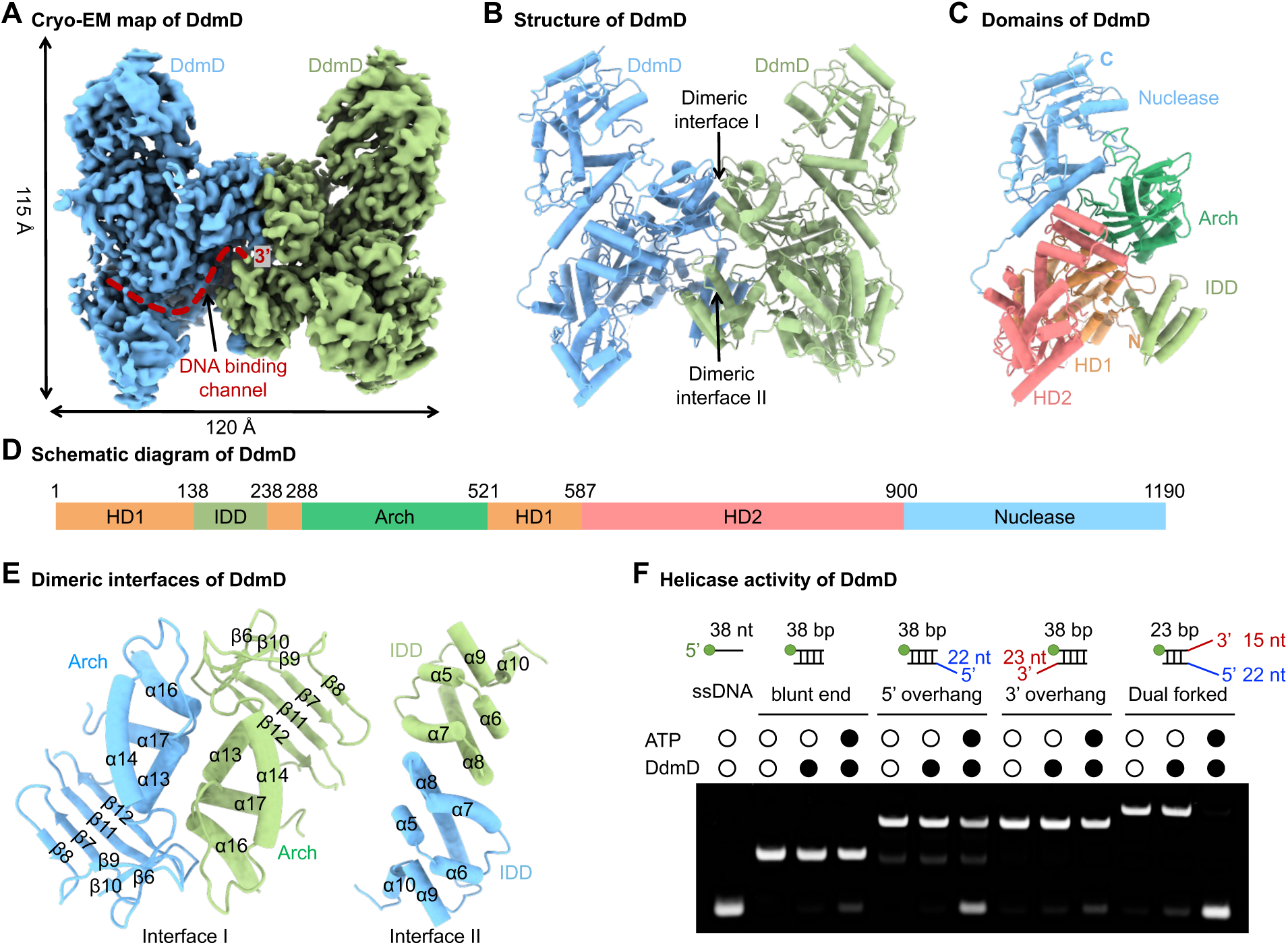
DdmD forms a dimer with helicase activity. (A) Cryo-EM density map of apo DdmD with the two protomers colored in blue and green, respectively. (B) Cylinder representation of apo DdmD. (C) Cylinder representation of a DdmD protomer, with each domain distinctly colored. (D) Schematic domain organization of DdmD, with residue numbers indicated above. (E) Cylinder representation of DdmD dimerization interfaces I and II, with secondary structures labeled. (F) The helicase activity of DdmD against various substrates with or without ATP. Green dots indicate 5′ Cy3 label. Substates are detailed in Table S2.

HD1 and HD2, two canonical RecA-like helicase domains common to all the SF2 helicases, form the helicase core of DdmD and create an ATPase active site at their interface. These domains contain seven conserved motifs (Ia & I to VI) of the SF2 helicase family (Figure S1D). ATP binds to the central cleft between HD1 and HD2, coordinated by the canonical motifs I, II, III, V, and VI ^12^(Figure S1D). Positioned above this central cleft is the Arch domain, analogous to that in other SF2 helicases like DinG ^21^ (Figure 1C). Unlike DinG’s Arch domain that consists of a four-stranded β sheet and four α-helices, DdmD’s Arch domain adopts a “hotdog”-like fold with a central long α-helix in the middle surrounded by three- and five-stranded β sheets and four α-helices (Figure S1E). Additionally, HD1 features an inserted domain (IDD) that resembles the iron-sulfur cluster domain of DinG. However, no iron-sulfur cluster is present in the DdmD IDD domain (Figures 1C and 1D). Together, the HD1, IDD, HD2, and the Arch domain form a typical SF2 helicase.

Above these helicase domains sits the DdmD nuclease domain, a member of the PD- (D/E)XK family^15^. It contains a conserved catalytic site composed of residues E1059, E1082, D1085, D1100, and K1102 (Figure S1F). The core structure of the nuclease domain adopts a conserved α-β-α fold, featuring a mixed β-sheet flanked by α-helices on both sides (Figure S1F). Furthermore, the nuclease domain includes a flower-mop- shaped region composed of two pairs of α-helical hairpins and one anti-paralleled β-sheet (Figure S1F), serving to anchor and connect the core domain with the HD2 domain of DdmD.

DdmD dimerization is governed by two interfaces: Interface I and Interface II, mediated by its Arch domain and IDD respectively, with a combined buried surface area of ∼5,100 Å^2^ (Figures 1B and 1E). In Interface I, the α-helical bundles of the Arch domain pack against each other, facilitating DdmD dimerization through extensive charge-charge and hydrophilic interactions (Figures 1E and S1G). In Interface II, on the other hand, both hydrophobic and hydrophilic residues in the IDD contribute to the dimerization (Figures 1E and S1H). In comparison to DinG, the dimeric arrangement of DdmD positions the 3’ end of the typical DNA binding channel in opposite orientations at the dimeric interface, potentially occluding the DNA exit sites and representing an autoinhibited state (Figure 1A).

Together, DdmD assembles as a dimer with two interfaces mediated by the Arch and IDD domains, potentially representing an autoinhibited state.

### DdmD unwinds DNA in the presence of ATP

To determine whether the dimeric DdmD has helicase activity, we tested different substrates, including blunt-ended double-stranded DNA (dsDNA), dsDNA with a 3’ overhang or a 5’ overhang, and dual-forked dsDNA, in the presence and absence of ATP (Figure 1F). We found that DdmD displayed robust helicase activity towards dual-forked dsDNA while showing weak activity towards dsDNA with a 5’ overhang (Figure 1F). In contrast, DdmD cannot unwind blunt-ended dsDNA or dsDNA with a 3’ overhang (Figure 1F). Moreover, DdmD helicase activity depends on ATP, akin to DinG. Consistently, an E273A mutant, which disrupts ATP hydrolysis, substantially reduced the helicase activity of DdmD (Figure S1I).

To further explore whether DNA substrates can regulate the ATP hydrolysis of DdmD, we tested the ATPase activity of DdmD in the presence of various DNA substrates. We found that both ssDNA and dsDNA can considerably enhance the ATP hydrolysis of DdmD, indicating direct interactions between various DNA substates with DdmD (Figure S1J). Further binding assays showed that DdmD indeed can preferably bind to ssDNA (Figure S1K).

Together, DdmD dimer displays robust helicase activity towards dual-forked dsDNA in an ATP-dependent manner and its ATPase activity is enhanced by binding to DNA substrates.

### DdmD and DdmE form a complex in the presence of DNA

To examine whether DdmD and DdmE form a complex for anti-plasmid defense, we biochemically reconstituted both proteins. Unexpectedly, purified DdmD and DdmE did not form a stable complex upon incubation (Figure 2A). Similarly, co-expression in *E. coli* did not yield a stable DdmD-DdmE complex (Figure S2A). These results led us to speculate that DdmD and DdmE might interact upon binding to DNA substates. After testing various DNA substrates, we discovered that a dual-forked DNA substrate, termed EHJ12, facilitates the formation of a stable complex suitable for subsequent structural analysis (Figures 2A and S2B-S2E).

**Figure 2.**
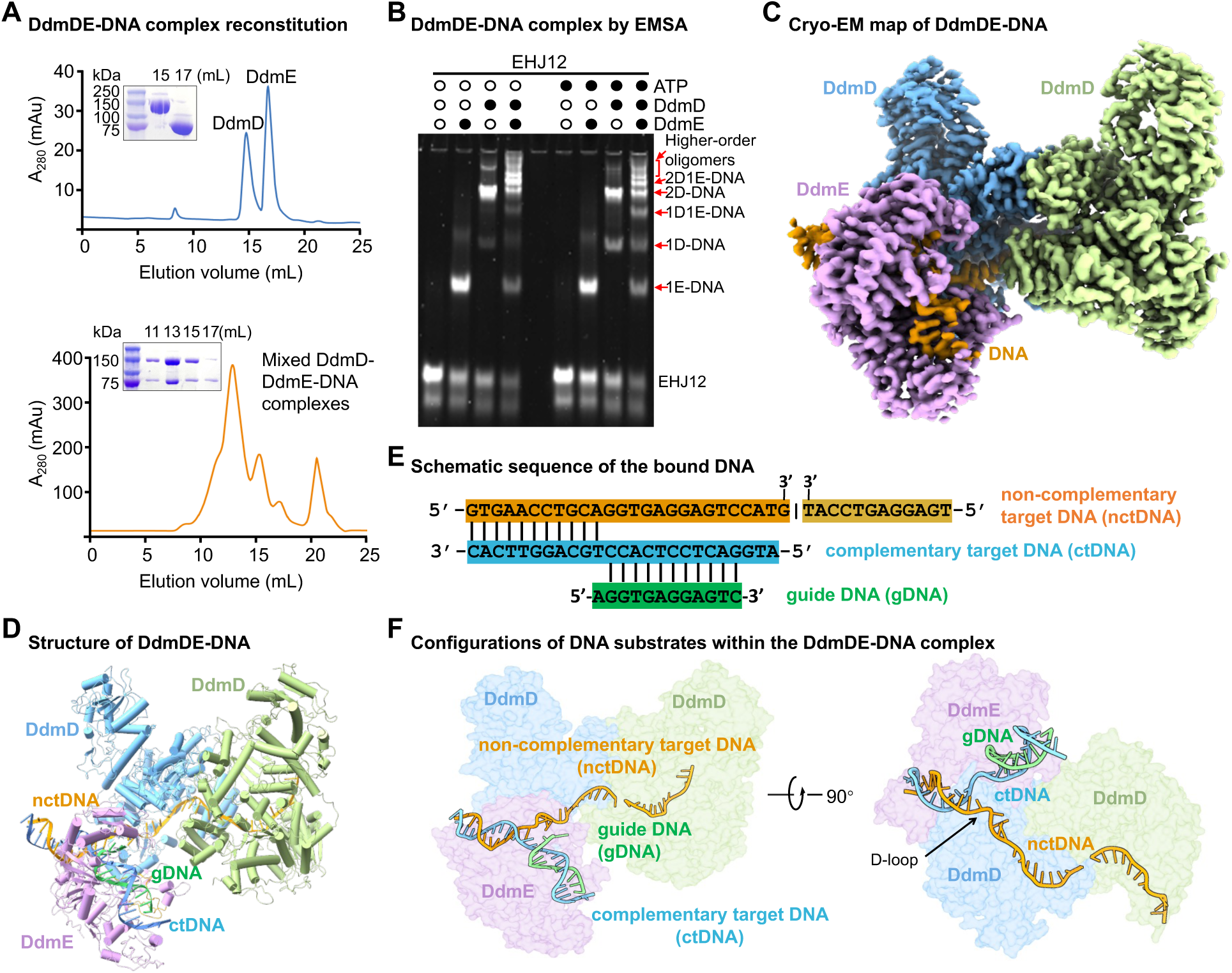
Structure of DdmD-DdmE-DNA ternary complex. (A) Gel filtration profiles and SDS-PAGE analyses of DdmD and DdmE after co-incubation without (top panel) or with (bottom panel) a dual-forked DNA substrate, EHJ12. (B) Electrophoretic Mobility Shift Assay (EMSA) analysis of the DdmD dimer and/or DdmE in complex with DNA substrates (EHJ12), with or without ATP. 1E, DdmE monomer; 1D, DdmD monomer; 2D, DdmD dimer. (C) Cryo-EM density map of DdmD-DdmE-DNA (2D-1E-DNA) ternary complex. (D) Schematic sequence of the visible DNA within the DdmD-DdmE-DNA ternary complex. (E) Ribbon representation of DdmD-DdmE-DNA ternary complex. (F) Ribbon representation of DNA substrate within the DdmD-DdmE-DNA ternary complex. The three DNA strands are designated as guide DNA (gDNA), complementary target DNA (ctDNA), and non-complementary target DNA (nctDNA).

A broad peak for the complex was observed in gel filtration, prompting us to run a native-PAGE to verify the complex’s identity (Figures 2A, S3A, and S3B). We identified several species, including DdmD-DdmE-DNA ternary complexes with different stoichiometries for DdmD and DdmE, the complex of DdmD dimer with DNA, and the complex of DdmD monomer with DNA, and the complex of DdmE with DNA (Figures S3A and S3B). Further electrophoretic mobility shift assays (EMSA) revealed that the incubation of DdmD, DdmE, and the EHJ12 DNA substrate led to the formation of the aforementioned complexes (Figure 2B). Notably, the addition of ATP during the incubation substantially promotes the generation of the monomeric DdmD-DNA complex, indicating that ATP may facilitate the disassembly of the DdmD dimer upon DNA binding (Figure 2B). Additionally, higher-order oligomers of the DdmD-DdmE-DNA ternary complex were observed both on gel filtration and in the EMSA experiment, indicating that multiple copies of DdmD and DdmE can simultaneously target to a DNA substrate for anti-plasmid defense (Figures 2B, S3A, and S3B).

To elucidate the assembly mechanism of the DdmD-DdmE-DNA complex, we determined cryo-EM structures of the reconstituted complexes eluted from gel filtration (Figures 2A, S2C-S2E, and Table S1). Consistently with our biochemical analysis, we obtained cryo-EM maps for a few different complexes. Among them, the best cryo-EM map for the DdmD-DdmE-DNA ternary complexes yielded a nominal resolution of 3.28 Å (Figures S2C-S2E, S3C, and Table S1). The structure revealed that the DNA substrate connected one DdmE monomer and one DdmD dimer tightly (Figures 2C-2E). The asymmetric assembly of the ternary complex indicates that one DdmE molecule can recruit one copy of DdmD dimer to the DNA substrate (Figure 2C).

Within the ternary complex, the DNA substrate has three strands with a markedly distorted configuration, indicating that DdmDE recognizes and targets DNA substrates via a DNA-guided DNA targeting mechanism (Figures 2E and 2F). As such, we designated these DNA strands as guide DNA (gDNA), complementary target DNA (ctDNA), and non-complementary target DNA (nctDNA) (Figures 2E and 2F). DdmE binds to the ctDNA in a guide-dependent manner, leading to the formation of a partial D-loop on the target DNA (Figure 2F). The 5’-end of the nctDNA and ctDNA form a short duplex captured by DdmE. Immediately following that duplex is a kinked nctDNA region in a single stranded form, extending all the way to the DNA binding channel of one DdmD protomer that directly interacts with DdmE (Figure 2F). In contrast, the other DdmD protomer was observed to bind a short piece of single strand DNA (Figure 2F). These data indicate that the two protomers of the DdmD dimer may bind to the ctDNA and nctDNA strands, respectively, upon D-loop formation under physiological conditions. Additionally, our biochemical analysis revealed that a short DNA fragment was generated from the original long DNA substrates when incubated with DdmDE, potentially serving as the guide DNA for DdmE (Figure S3D).

Together, our biochemical and structural analyses indicated that one DdmE molecule recruits and loads one DdmD dimer to the target DNA, potentially via a DNA-guided DNA-targeting mechanism.

### The DdmDE complex eliminates plasmids

DdmE exhibits a typical Argonaute domain architecture similar to other pAgos, comprising N, L1, PAZ, L2, and MID domains, followed by a PIWI-like domain with a central DNA binding groove^16,23,24^ (Figures 3A-3C). In comparison to *Cb*Ago^24^, the central DNA-binding groove of DdmE is wider (Figure 3C). Unlike the canonical PAZ domain, the PAZ domain of DdmE is notably small, consisting of about 40 residues forming a hairpin-like structure (Figures 3A and 3B). Remarkably, the PIWI domain includes an inserted domain responsible for interacting with DdmD, which we termed the DdmDE-interacting domain (DID) (Figures 3A and 3B). The DID is composed of four α-helices, forming a hairpin bodkin-like structure without a known homolog in the PDB database, suggesting that it may represent a new fold (Figures 3B and 3D). Both the N-terminal and C-terminal ends of DID are connected to the PIWI-like domain by long loops, positioning the DID spatially distant from the PIWI-like domain (Figure 3D). The DID is located adjacent to the L1-PAZ- L2 domains, extending beyond the core Argonaute fold to form a pocket with the L1-PAZ- L2 on the opposite side of DdmE’s DNA binding groove (Figures 3E and 3F).

**Figure 3.**
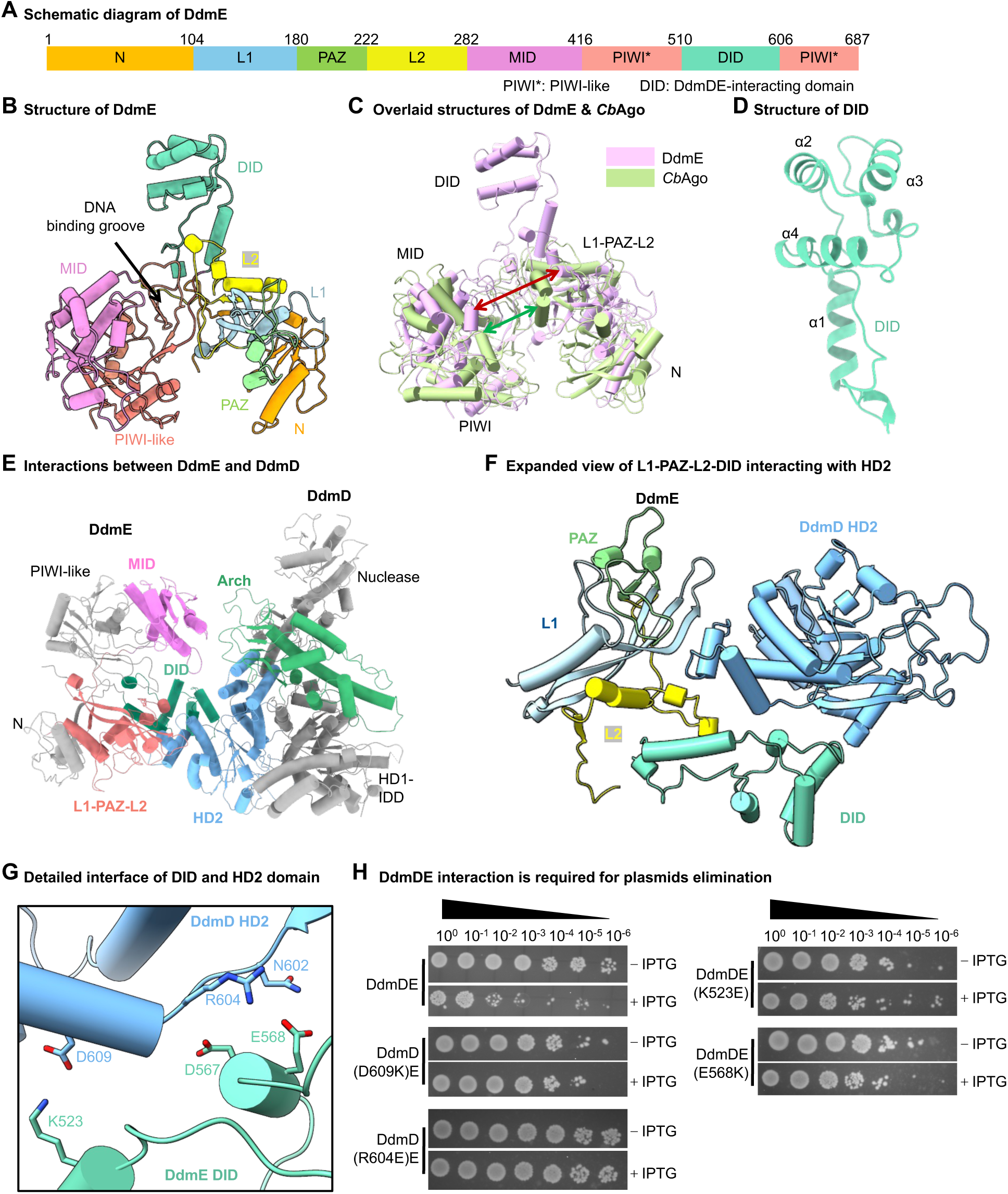
DdmE rectuits DdmD for eliminating plasmids. (A) Schematic domain organization of DdmE with residue numbers indicated above. (B) Cylinder representation of DdmE, with each domain distinctly colored as shown in (A). (C) Overlaid structures of DdmE and CbAgo reveal the DdmDE-interacting domain (DID) in DdmE. (D) Ribbon diagram of DdmE DID domain with secondary structures labeled. (E) Overview of DdmE and its engagement with one DdmD protomer, depicted in cylinder representation with interacted domain distinctly colored and labeled. (F) Interaction interfaces of DdmE and DdmD mediated by L1-PAZ-L2-DID and HD2 domains, with each domain distinctly colored and labeled in cylinder representation. (G) Detailed interactions between DdmE DID domain (pale green) and DdmD HD2 domain (sky blue). (H) Bacterial growth assay showing interactions between DdmD and DdmE are critical for plasmids elimination.

DdmD interacts with DdmE using its HD2 and Arch domains. An α-helical hairpin in DdmD’s HD2 nests within the pocket formed by DdmE’s DID and L1-PAZ-L2 domains, primarily through hydrophilic interactions (Figures 3F, 3G, and S3E). Specifically, DdmD’s HD2 and DdmE’s DID engage in hydrogen-bonding and charge-charge interactions, including salt bridges between D609 of DdmD and K523 of DdmE, and between R604 of DdmD and E568 of DdmE (Figure 3G). Additionally, residues from the L1-PAZ-L2 domains also contribute to both charge-charge interaction and hydrogen bonding with DdmD (Figure S3E). Weak interactions are observed between the Arch domain of DdmD and the MID domain of DdmE through hydrogen-bonding (Figure S3F). Together, DdmD and DdmE form an electrostatic and hydrophilic interface with a buried surface area of ∼800 Å^2^, consistent with previous biochemical reconstitution showing their weak interactions in the absence of DNA substrates (Figure 2A).

To confirm the functional importance of the DdmDE interaction, we developed an *in vivo* bacterial growth assay (Figure S3G). Upon induction, *E. coli* cells co-expressing DdmD and DdmE lost their plasmids and failed to grow on antibiotic-containing plates (Figure S3H). In contrast, interfacial mutants of DdmDE partially or fully rescued *E. coli* growth on these plates (Figure 3H), underscoring the critical role of the DdmDE interaction in anti-plasmid defense.

Together, DdmE, an pAgo-like protein, directly interacts with DdmD to facilitate anti-plasmid defense.

### DdmE is a DNA-guided DNA-targeting module

DdmE exhibits distinct features in coordinating its DNA substrate compared to other well-characterized Argonautes (Figures 4A-4C, and S4A-S4C). First, the DNA substrate of DdmE consists of three strands, gDNA, ctDNA, and nctDNA, contrasting with the two-stranded substrates seen in other Argonautes^16^ (Figures 4A and 4B).

**Figure 4.**
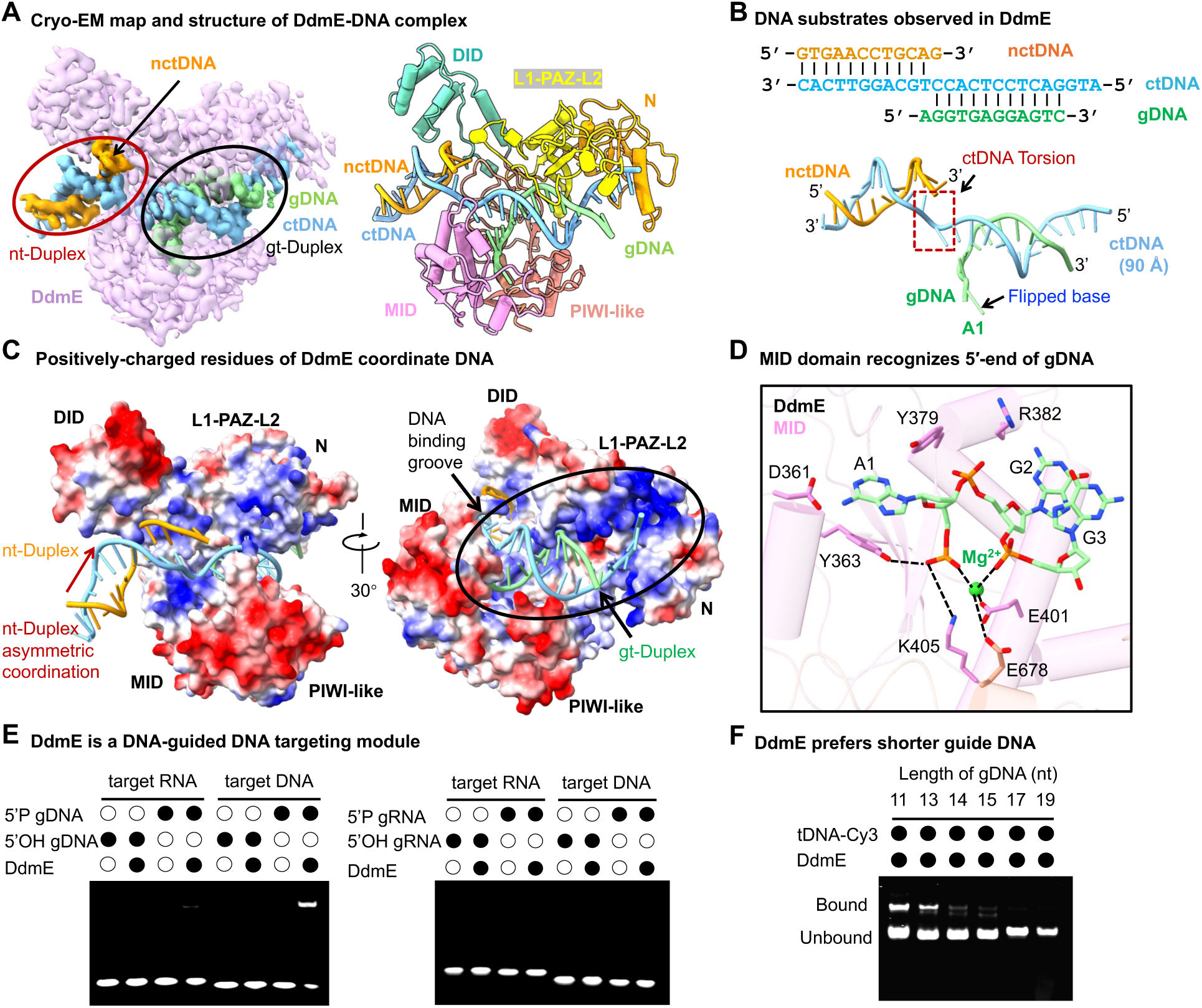
DdmE is a DNA-guide DNA-targeting module. (A) Cryo-EM density map and cylinder representation of DdmE and DNA complex. gDNA, guide DNA; ctDNA, complementary target DNA; and nctDNA, non-complementary target DNA. Nt-Duplex and gt-Duplex are higlighed in red and blue circles, respectively. (B) Schematic sequence and structure of gDNA, ctDNA, and nctDNA in DdmE. (C) DNA coordination by DdmE. DdmE is shown as electrostatic surface potential model. nt-Duplex and gt-Duplex interact with DdmE differently. (D) Expanded view of the 5’ nucleotide of guide DNA coordinated by residues in DdmE MID domain. (E) DdmE uses 5’-phosphorylated gDNA for DNA targeting, revealed by EMSA. Target DNA and RNA are labeled by Cy-3 fluorophore. (F) DdmE prefers gDNA shorter than 14 nt, revealed by EMSA.

Second, the extended ctDNA, approximately 90 Å in length, spans the entire DdmE molecule (Figures 4A and 4B). Its 5’-end forms a duplex with gDNA (gt-Duplex) while its 3’-end forms a duplex with nctDNA (nt-Duplex) (Figure 4A). In contrast, other Argonautes bind only a single duplex.

Third, the ctDNA displayed a distorted configuration due to its interactions with DdmE (Figures 4B, S4A, and S4C). Part of the ctDNA lies in DdmE’s central groove, while the other part interacts with residues in the DID domain (Figures 4C and S4C). In the central groove, ctDNA and gDNA pair to form the gt-Duplex, coordinated by charged residues on both sides (Figure 4C). Conversely, the remaining part of ctDNA forms the nt-Duplex, engaging the DID by only one side (Figure 4C). This asymmetric interaction tilts the nt- Duplex towards DID, partially explaining the distorted configuration of ctDNA (Figure 4C). Furthermore, several bulky residues of DdmE, including K625, R664, H393, and H663, form a narrow neck, interacting with the distorted region of ctDNA and stabilizing its configuration (Figure S4A).

Fourth, the 5’-end of ctDNA is capped by DdmE’s N domain (Figure 4C). Consequently, 14 nucleotides (nt) of ctDNA were observed within DdmE’s central groove, pairing with a 12 nt gDNA in our structure (Figure S4C). This suggests that DdmE may prefer a gDNA shorter than 14 nucleotides, as longer guides may cause steric clashes with DdmE’s N domain.

Fifth, residues K230 and R232 within a loop of the L2 domain penetrate the minor groove of gtDNA, likely serving as a sensor for B-form duplex and contributing to DdmE’s specificity for DNA (Figure S4B).

The MID domain of DdmE, similar to those in other Argonautes, is responsible for recognizing the 5’-end of gDNA (Figures 4D and S4D). The first nucleotide at the 5’-end of gDNA is flipped out from the duplex and captured by a hydrophobic pocket composed of K405, Y363, and Y379, which has not been seen in any other Argonaute proteins^16,25,26^ (Figures 4D and S4D). The phosphate group of the 5’-end gDNA is coordinated by specific residues, indicating a preference for 5’-phosphorylated gDNA (Figure 4D). Moreover, the proximity of a magnesium ion to the phosphate of the 5’-end gDNA underscores DdmE’s magnesium-dependent recognition of gDNA, common in prokaryotic Argonaute proteins ^8,25,27^ (Figures 4D and S4D). Additional residues in the pocket, including E401 and E678, are involved in coordinating the magnesium ion (Figure 4D).

Our structural analysis strongly suggests that DdmE recognizes DNA substrate via a DNA-guided DNA-targeting mechanism (Figures 4A and 4B). To test this hypothesis, we conducted EMSA assays to evaluate DdmE’s substrate specificity. DdmE effectively binds to target DNA in the presence of 5’-phosphorylated gDNA, but not gDNA with a 5’-hydroxyl (5’-OH) group (Figure 4E). DdmE also does not bind to target RNA in the presence of gDNA (Figure 4E). Furthermore, DdmE shows no affinity for target DNA or target RNA in the presence of 5’-phosphorylated gRNA or 5’-OH gRNA (Figure 4E). Interestingly, DdmE effectively binds to 5’-Cy3 modified DNA *in vitro* (Figure S4E), potentially due to extra space around the 5’-monophosphate-binding site, a feature resulting from DdmE’s incomplete PIWI-helical subdomain compared to typical Argonautes^16^. Further EMSA tests revealed that DdmE prefers guides shorter than 14 nt, with the strongest binding affinity observed for 11 nt, consistent with our structural analysis (Figure 4F). Among gDNAs ranging from 11 to 19 nt, shorter guides were more potent (Figure 4F).

DdmE lacks nuclease activity towards target DNA in the presence of gDNA or towards other DNA substrates, consistent with the absence of the canonical DEDX catalytic tetrad in its PWI-like domain^28^ (Figures S4F and S4G). These findings support the classification of DdmE as a catalytically inactive Argonaute-like protein.

Together, DdmE is a DNA-guided DNA-targeting module with a preference for 5’- phosphorylated gDNA shorter than 14 nt.

### DdmD recognizes ssDNA

Upon binding to ctDNA in a guide-dependent manner, DdmE recruits and loads the DdmD dimer onto the target DNA (Figure 2F). In the DdmD-DdmE-DNA ternary complex, both protomers of DdmD uses their helicase domains to bind to ssDNA, similar to other SF2 family helicases ^21^ (Figure 2F). We further determined the cryo-EM structure of the DdmD dimer in complex with DNA, revealing a conserved mechanism for DdmD’s recognition of DNA (Figures 5A, 2F, and S2C-S2E). A kinked ssDNA with flipped bases extends across the HD2 domain, traverses to the HD1 domain, and terminates at the dimeric interface of DdmD (Figures 5A-5F). Four bases are coordinated by the HD1 domain, five interact with the HD2 domain, one base in the middle of the HD1 and HD2 domains, and one at the dimeric interface (Figures 5D-5F). We termed the first nucleotide of the nctDNA strand, where the ctDNA enters the HD2 domain of DdmD and forms a kink, as G1 (Figure 5E). Following G1, nucleotides A2 and G3 exhibit a flipped conformation and form a continuous stack with G1, a feature not observed in other structurally characterized SF2 helicases like DinG^21^(Figure 5E and S5A). At nucleotide G3, the nctDNA makes a second kink as it packs against F639 via a χ-χ interaction and passes through a restriction point formed by Q781, T780, and R669 (Figure 5E). Structural analysis reveals that the HD2 domain of DdmD possesses an α-helical bundle comprising three α-helices at the ssDNA entrance (Figures 5E and S5B). This bundle acts as an obstacle, distorting the first three bases of ssDNA (Figures 5E and S5C).

**Figure 5.**
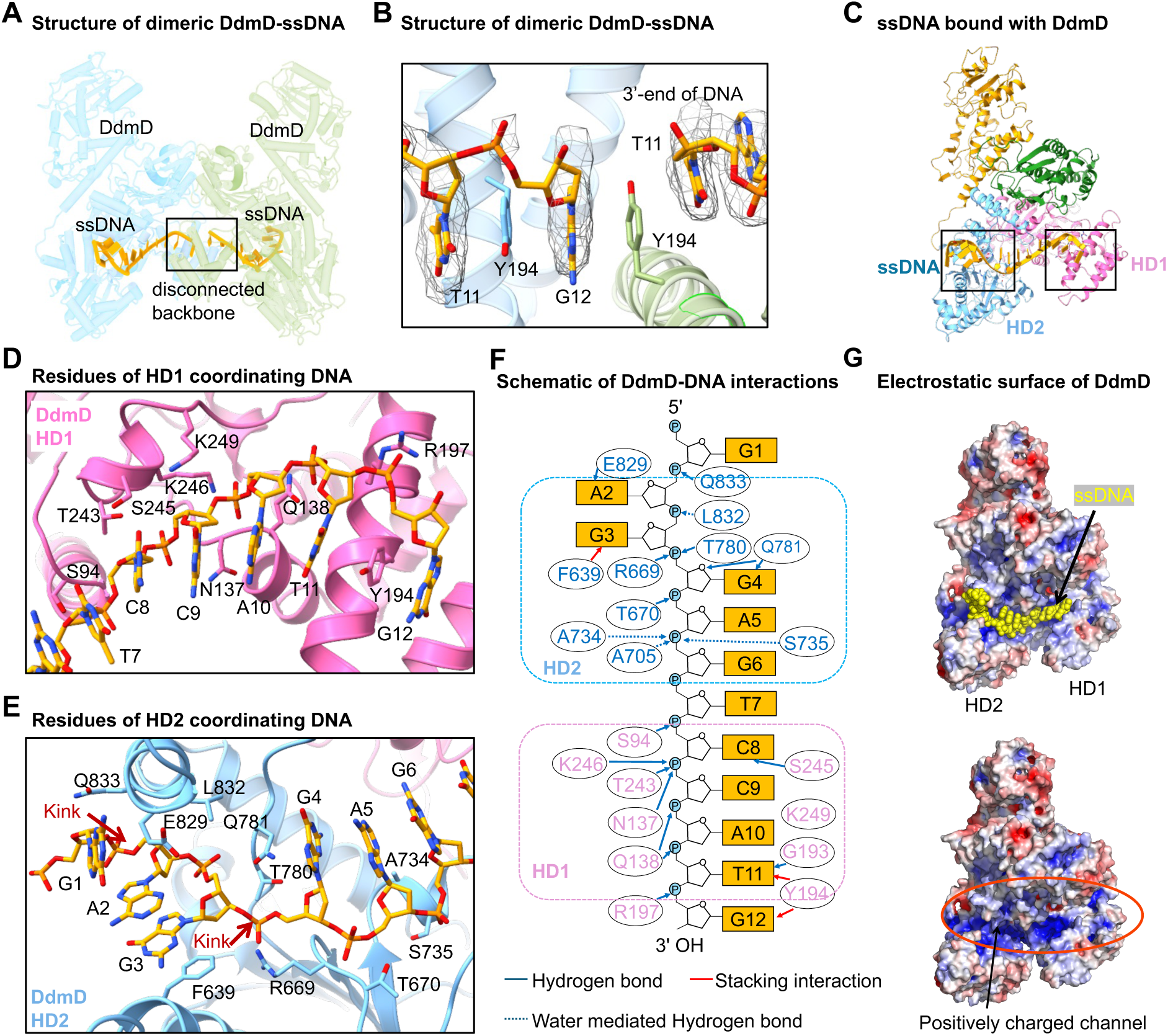
DdmD recognizes ssDNA. (A) Cylinder representation of DdmD with single strand DNA (ssDNA). (B) A close-up view of the disconnected backbone, as indicated in (A). (C) Ribbon diagram of DdmD with ssDNA. ssDNA, HD1 domain and HD2 domain of DdmD were distinctly colored as indicated. (D) Detailed view of ssDNA coordinated by key residues in DdmD HD1 domain. (E) Detailed view of ssDNA coordinated by key residues in DdmD HD2 domain. (F) Schematic view of residues of DdmD coordinating ssDNA. (G) Representation of the positively charged channel within DdmD HD1 and HD2 domains, crucial for ssDNA binding.

The bound DNA occupies a positively charged channel across the helicase domain, following a 5’ to 3’ polarity, akin to some SF2 helicases such as DinG ^17,21^ (Figures 5G and S5A). Interactions between ssDNA and DdmD are predominantly mediated by the phosphodiester backbone and hydrophilic residues within DdmD’s channel (Figures 5F and 5G). The polarity of DdmD-bound DNA suggests an unwinding mechanism shared by DdmD and DinG^21^, indicating that DdmD translocates along DNA with a 5’ to 3’ direction. Both HD1 and HD2 harbor a positively charged patch, likely alternating to securely grip the ssDNA during DNA unwinding and translocation (Figure 5G). Remarkably, the 3’-end guanine of ssDNA is terminated and stabilized by Y194 through π-π interactions, halting further extension of ssDNA within DdmD’s channel (Figure 5B). Our structural analysis is consistent with the fact that DdmD preferred to bind DNA substrates with a 3’-overhang over a 5’-overhang or blunt-end dsDNA in our EMSA assays (Figure S5D).

Together, DdmD binds to ssDNA and translocates along DNA with a 5’ to 3’ polarity.

### DNA disassembles the autoinhibited DdmD dimer

To understand the impact of nucleic acid binding on DdmD, we compared the dimeric interfaces of DdmD in its apo state and ssDNA-bound state. Upon DNA binding, large conformational changes occurred at the dimeric interface, causing the DdmD protomers to tilt away from each other and reducing their interactions (Figure 6A). Consequently, the total buried area of the DdmD dimer in the DdmD-DNA complex is approximately 4,700 Å^2^, smaller than that of the apo DdmD dimer (∼5,100 Å^2^), indicating that DNA binding indeed weakened the interactions between the two protomers (Figure 6A). Furthermore, several key charged interactions on both interface I and Interface II are either reduced or completely abolished in the DNA bound DdmD dimer (Figures 6B and S6A). Our structural analysis indicated that DNA binding promotes DdmD dimer disassembly by reducing interactions between the two protomers.

**Figure 6.**
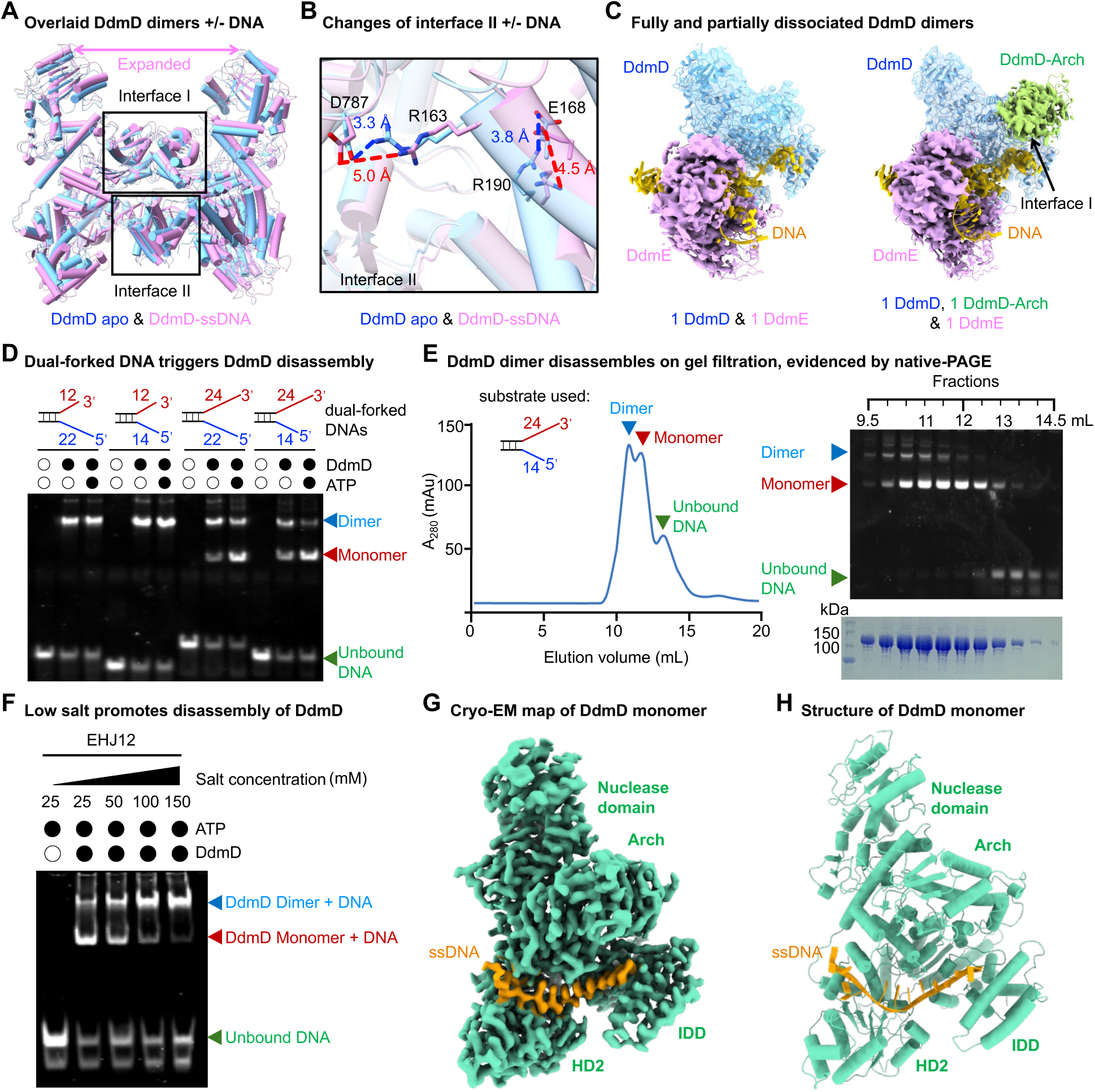
Forked DNA triggers disassembly of DdmD dimer. (A) Overlaid structures of apo DdmD (cyan) and dimeric ssDNA-bound DdmD (pink). (B) Comparison of key residues of interface II in apo and DNA-bound DdmD dimers. (C) Cryo-EM density maps of fully dissociated and partially dissociated DdmD-DdmE- DNA (1D-1E-DNA) ternary complexes. (D) Forked DNA with a long 3’ overhang triggers the DdmD dimer dissociation, revealed by EMSA. The lengths of the overhang are indicated. All the DNA substrates are detailed in Table S2. (E) Forked DNA with a long 3’overhang triggers the dissociation of the DdmD dimer, revealed by gel filtration and native-PAGE. (F) Low salt conditions promote DdmD disassembly, revealed by EMSA. (G) Cryo-EM density map of ssDNA-bound DdmD monomer. (H) Cylinder representation of ssDNA-bound DdmD monomer.

To investigate whether DNA binding could effectively trigger the disassembly of the DdmD dimer, we conducted a deep classification of the DdmD-DdmE-DNA complex, resulting in two additional maps: one depicting a single copy of DdmD and the other showing partial density for the second copy of DdmD (Figures 6C, S2D, and S2E). These maps represent a fully dissociated state and a partially dissociated state during the disassembly of DdmD triggered by DNA binding. In the fully dissociated state, one copy of the DdmD monomer was observed to form a complex with DdmE and DNA (Figure 6C). In the partially dissociated state, Interface II has been completely disrupted while Interface I remains intact (Figure 6C). As such, the cryo-EM density corresponding to the Arch domain in the neighboring DdmD is visible (Figure 6C), indicating that the disruption of Interface II may be the initial step in DdmD dimer disassembly. This is consistent with the observation that DNA binding affects Interface II more severely than Interface I (Figures 6B and S6A). This analysis suggested that DdmD disassembly, triggered by DNA binding, may involve in two steps with Interface II being disrupted first.

To further support that DNA binding can trigger the disassembly of DdmD, we incubated DdmD with various DNA substrates and analyzed these samples by EMSA, gel filtration, native polyacrylamide gel electrophoresis (native-PAGE), and structural analysis (Figures 6D, 6E, and S6B-S6D). Our EMSA assays showed that a dual forked DNA with longer unpaired nucleotides effectively triggered the dissociation of the DdmD dimer, while a dual forked DNA with shorter unpaired nucleotides did not (Figure 6D). Further analysis revealed that the length of 3’-overhang plays a critical role in triggering the disassembly of the DdmD dimer (Figures 6D and S6E). Dual-forked DNA substrates with a 3’-overhang longer than 18 nt effectively triggered the DdmD dimer disassembly, whereas those with a 3’-overhang shorter than 14 nt did not (Figure S6E). Additionally, ATP can promote the disassembly of DdmD, suggesting that helicase activity may be coupled with the disassembly of DdmD (Figures 6D and S6E). Our EMSA findings were further confirmed through gel filtration and native PAGE analysis, supporting that dual-forked DNA with a longer 3’-overhang can trigger the assembly of the DdmD dimer (Figure 6E). Moreover, low salt conditions facilitate the disassembly of DdmD, consistent with the hydrophobic nature of DdmD dimeric Interface II being disrupted first during disassembly (Figures 6C, 6F, and S1H).

We further determined the cryo-EM structures of DdmD incubated with a dual-forked DNA, a sample used for our gel filtration analysis (Figures 6E and S6B). The selected 2D class averages consisted of more than 80% DdmD monomers, with less than 20% DdmD dimers, consistent with our gel filtration analysis that the dual-forked DNA disassembles the DdmD dimer (Figures 6E, S6C, and S6D). The disassembled DdmD monomer was reconstructed to a resolution of ∼3.0 Å, with the bound ssDNA clearly resolved (Figures 6G, 6H, and S6B). Compared with the ssDNA bound dimer, the DdmD monomer exhibits subtle but visible conformational changes in the Arch and IDD domains, which are critical for DdmD dimerization (Figures 1E and S6F). In both domains of the DdmD monomer, key α-helical bundles for dimerization tilt towards the direction where dimeric interfaces occur in the DdmD dimer, providing the structural basis of DdmD disassembly (Figure S6F).

Together, these findings suggest that the DdmD dimer disassembles into monomers upon binding to forked DNA substrates, with interface II being disrupted first, followed by interface I.

### DdmD translocates along and eliminates plasmids

DdmD contains a nuclease domain homologous to PD-(DE)XK nucleases, potentially harboring one metal-catalysis mechanism^15,29^. When we incubated DdmD with ssDNA in the presence of various metal ions, we found that DdmD displayed robust nuclease activity with manganese, modest activity with magnesium, and no activity with calcium, nickel, or zinc (Figure 7A). Substituting alanine for residues D1059 and H1106 abolished the nuclease activity of DdmD, highlighting their crucial roles in catalysis (Figures 7A and S1F). Further experiments with various substrates revealed that DdmD could cleave all DNA substrates containing an ssDNA fragment but could not cleave blunt-end dsDNA (Figures 7B and S7A). This finding aligns with our EMSA assay that DdmD requires a single-stranded portion to recognize DNA substrates (Figures 7B, S7A, and S5D). Additionally, in the presence of magnesium, ssDNA can be gradually degraded by DdmD with low efficiency (Figure S7B). These findings suggested that the DdmD dimer can cleave ssDNA in the presence of magnesium or manganese.

**Figure 7.**
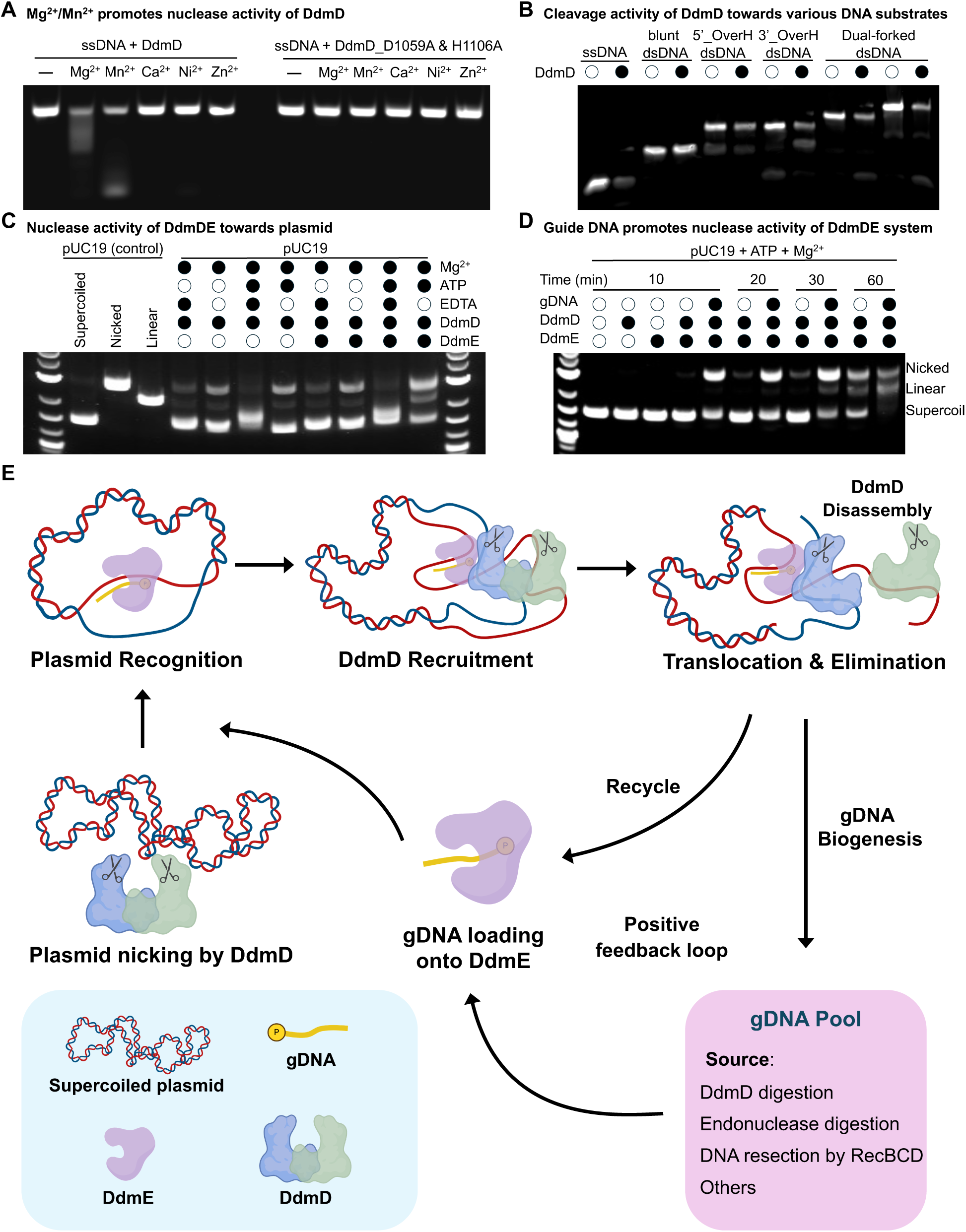
DdmDE eliminates plasmids by DNA-guided DNA targeting. (A) DdmD nuclease activity with various divalent metal ions. DdmD_D1059A_H1106A is a catalytic inactive mutant. (B) DdmD nuclease activity against various DNA substrates in the presence of Mn^2+^. DNA substrates are detailed in Table S2. (C) DdmDE nuclease activity against pUC19 *in vitro*. Nb.BsrDI was used to generate nicking plasmid as control. EcoRI was used to generate linear plasmid. (D) gDNA enhance nuclease activities of DdmDE, but not DdmD alone. (E) Mechanism for the DdmDE-mediated plasmid elimination.

To investigate whether DdmD and DdmDE can eliminate plasmids, we developed an *in vitro* plasmid elimination assay. Incubating DdmD alone with plasmids showed that DdmD produced nicked open circular plasmid for pUC19 (a small plasmid), and both linear and circular plasmids for pBAD (a large plasmid) in the presence of magnesium (Figures 7C and S7C). The addition of EDTA abolished DdmD’s activity, further underscoring the importance of magnesium in catalysis. ATP had little effect on the plasmid cleavage by DdmD (Figures 7C and S7C). In contrast, the DdmDE complex displayed much higher nuclease activities towards both pUC19 and pBAD in the presence of ATP, converting the majority of supercoiled plasmids into linear and circular forms (Figures 7C and S7C). This indicates that the helicase activity of DdmD may enhance the nuclease activity of DdmD only in the presence of DdmE. Moreover, in the presence of guide DNA, the DdmDE, but not DdmD alone, developed substantially higher nuclease activity towards pUC19 plasmids, highlighting the functional importance of gDNA in plasmid elimination by DdmDE (Figure 7D).

Together, these results indicates that DdmD alone has residual nuclease activity and may relax supercoiled DNA in bacteria to facilitate DdmE targeting. In the presence of DdmE and gDNA, the DdmD can be effectively recruited onto target plasmids for disassembly, translocation, and cleavage.

## DISCUSSION

### Mechanism of DdmDE anti-plasmid defense

Here, we provide mechanistic insights into plasmid elimination by the DdmDE system, a recently identified anti-plasmid defense system in *Vibrio cholerae* ^10^. We demonstrate that DdmDE effectively eliminates plasmids via a DNA-guided DNA-targeting mechanism (Figure 7E). DdmE, featuring a typical Argonaute fold with a DID domain for interacting with DdmD, binds a 5’-phosphorylated short ssDNA as a guide to target plasmid DNA. In its apo state, DdmD forms dimers with residual nuclease activity towards plasmids, resulting in nicked open circular plasmids (Figure 1 and Figure 7). These nicked open circular plasmids facilitate DdmE targeting (Figure 7E). Upon DdmE-gDNA binding to plasmids, a D-loop is formed to recruit DdmD (Figure 7E). Each DdmE molecule recruits a DdmD dimer onto plasmids, potentially with one DdmD protomer binding to the nctDNA strand and the other to the ctDNA strand. DNA binding induces conformational changes in DdmD, disrupting its dimerization interfaces (Figure 6). Our detailed biochemical and structural analyses show that forked DNA with a longer 3’ overhang can effectively trigger the transition of DdmD dimers into monomers (Figure 6). Consequently, this primes DdmD for the elimination of plasmids by ATP-driven translocation with a polarized direction of 5’ to 3’ (Figure 5). ATP enhances the plasmid eliminating activity of DdmDE through two mechanisms. First, the presence of ATP potentially enhances DdmD dissociation through binding-induced conformational changes in the IDD (Figure 6). Second, ATP can promote the translocation of disassembled DdmD along the plasmid for cleavage (Figure 1). Additionally, gDNA substantially enhances the plasmid degradation activity of DdmDE, but not DdmD alone, underscoring the importance of gDNA in plasmid targeting by DdmDE (Figure 7D).

The mechanism of DdmDE parallels that of the type I CRISPR-Cas system ^30,31^. In the type I CRISPR-Cas systems, Cascade, the RNA-guided effector complex, functions to recognize the target and recruit Cas3, a fusion protein of helicase and nuclease, to the non-target DNA strand^31^. Cas3 then cleaves the target DNA to generate short ssDNA fragments while translocating along the DNA using ATP, acting as an intact complex with Cascade^31^.

### Guide DNA recognition by DdmE

Structural analysis reveals that the MID-domain of DdmE preferentially recognizes the 5’- phosphate of the gDNA in a magnesium-dependent manner. Our EMSA assays confirmed that 5’-phoporylated gDNA enhances target DNA recognition, consistent with our structural findings. The first nucleotide at the 5’-end of gDNA, an adenosine, fits well into a pocket in the MID domain (Figure 4). This pocket in DdmE is large enough to accommodate all four types of nucleotides, indicating no sequence preference for the 5’- end of gDNA. Consistently, our binding assay showed effective binding of DdmE to gDNA with a Cy3 modification at the 5’-end (Figure S4). Given that DdmE interacts with the phosphodiester backbone of gDNA, it likely exhibits no sequence preference.

Despite no sequence preference to gDNA, DdmE shows a preference for the length of gDNA. Structural analysis indicated that longer DNA duplexes may clash with DdmE’s N-domain, suggesting a preference for shorter gDNA. Our EMSA assays supported this, showing that gDNA of 13 nt or shorter is more potent in DNA targeting (Figure 4F). Similarly, Jinek’s group reported that gDNA of 14 nt or shorter robustly promotes target DNA recognition by DdmDE^32^. Our EMSA results further showed an inverse correlation between gDNA length (11 nt to 19 nt) and targeting potency (Figure 4F).

### Dissociation, translocation and cleavage of DdmD

Our structural and biochemical analyses suggested that DdmD transitions from dimers to monomers upon DNA binding, enabling efficient translocation and cleavage (Figure 6). We propose that both DdmD protomers must bind DNA to trigger effective dissociation. Thus, DdmE likely recruits a dimeric DdmD to plasmid DNA, followed by DdmD dissociation of DdmD. The resulting DdmD monomers can efficiently translocate on plasmid DNA for cleavage, explaining why both hydrolase and nuclease activities of DdmD are required for plasmid elimination by DdmDE in bacteria^10^.

### Biogenesis of guide DNA

The guide DNA for DdmE can arise from various sources, including DdmD cleavage products, endonuclease-generated ssDNA fragments, and DNA end resection. Upon activation, DdmD cleaves plasmids, potentially producing small ssDNA fragments that act as guides for targeting the same plasmids. Indeed, during our reconstitution of the DdmD-DdmE-DNA complex, the guide DNA in DmDE likely originated from DdmD-processed substrates (Figure S3D), creating a positive feedback loop that enables more DdmE molecules to target the same plasmids. Additionally, bacterial endonucleases often generate DNA fragments with 5’-overhangs^33^, which DdmE can use as guides. DNA-end resection by the RecBCD complex also contributes to guide generation for other pAgos and CRISPR-Cas systems^9,34,35^. This dependence on RecBCD explains why DdmDE targets small multicopy plasmids lacking Chi sites. Other sources may also contribute to guide DNA generation.

In summary, our study defines DdmDE as a DNA-guided DNA-targeting system for plasmid elimination. From a translational standpoint, these intriguing features could potentially be harnessed as valuable tools in synthetic biology, such as exogenous DNA sensors.

## ACKNOWLEDGMENTS

We thank Dr. Wen Tang at OSU, Dr. Yuan He at Northwestern University, Dr. Shiyu Xia, at Caltech, and Anthony D. Rish at OSU for discussion and critical comments for the manuscript. We thank Dr. Melanie Blokesch from École Polytechnique Fédérale de Lausanne, Switzerland for providing the genomic DNA containing the DdmDE system. Grids screening was performed at OSU CEMAS with the assistance of Drs. Giovanna Grandinetti, Yoshie Narui, and Binbin Deng. Cryo-EM data were collected at PNCC and NCI cryo-EM national centers supported by grants from the NIH National Institute of Health Common Fund Transformative High Resolution Cryo-Electron Microscopy program. K.N. is supported by NIH R01GM124320 and R01GM138997. X.Y.Y. was supported by the center for RNA biology fellowship program of the Ohio State University.

## AUTHOR CONTRIBUTIONS

T.M.F. conceived the project. X.Y.Y. and Z. S. performed molecular cloning and biochemical purification and determined the cryo-EM structures. X.Y.Y., Z.S., and T.M.F. built the models. X.Y.Y. and S.Z. did all the biochemical assays and in vivo growth assay. T.M.F., Z.S., X.Y.Y., C.W., and N.K. analyzed the data. T.M.F. wrote the manuscript with inputs from all the authors.

